# Morphogenesis and morphometry of brain folding patterns across species

**DOI:** 10.1101/2025.03.05.641692

**Authors:** Sifan Yin, Chunzi Liu, Gary P. T. Choi, Yeonsu Jung, Katja Heuer, Roberto Toro, L. Mahadevan

**Author notes:** **For correspondence:** (LM).

## Abstract

Evolutionary adaptations associated with the formation of a folded cortex in many mammalian brains are thought to be a critical specialization associated with higher cognitive function. The dramatic surface expansion and highly convoluted folding of the cortex during early development is a theme with variations that suggest the need for a comparative study of cortical gyrification. Here, we use a combination of physical experiments using gels, computational morphogenesis, and geometric morphometrics to study the folding of brains across different species. Starting with magnetic resonance images of brains of a newborn ferret, a fetal macaque, and a fetal human, we construct two-layer physical gel brain models that swell superficially in a solvent, leading to folding patterns similar to those seen *in vivo*. We then adopt a three-dimensional continuum model based on differential growth to simulate cortical folding *in silico*. Finally, we deploy a comparative morphometric analysis of the *in vivo, in vitro*, and *in silico* surface buckling patterns across species. Our study shows that a simple mechanical instability driven by differential growth suffices to explain cortical folding and suggests that variations in the tangential growth and different initial geometries are sufficient to explain the differences in cortical folding across species.

## Introduction

Although not all brains are folded, in many mammals, the folded cerebral cortex is known to be critically important for brain cognitive performance and highly dependent on the hierarchical structure of its morphology, cytoarchitecture, and connectivity ***Gautam et al. (2015); Suárez et al. (2020); Pang et al. (2023***). Brain function is thus related both to the topological structure of neural networks ***Bullmore and Bassett (2011***), as well as the geometry and morphology of the convoluted cortex ***Kriegeskorte and Wei (2021***), both of which serve to enable and constrain neuronal dynamics ***Pang et al. (2023***). Across species, cortical morphologies show a large diversity, as shown in Fig. 1(a) ***Takahata et al. (2012); Heuer et al. (2019***). And within our own species and in model organisms such as the ferret used to study the genetic precursors of misfolding, cortical folding and misfolding are known to be markers of healthy and pathological neurodevelopment, disease and aging ***Hutton et al. (2009); de Moraes et al. (2024***) (see also (Fig. S1 ***Oegema et al. (2020); Choi et al. (2025***)). Thus a comparative study of cortical folding is essential for understanding brain morphogenesis and functionalization across evolution ***Pang et al. (2023); Schwartz et al. (2023***), during development as well as in pathological situations associated with disease.

**Figure 1.**
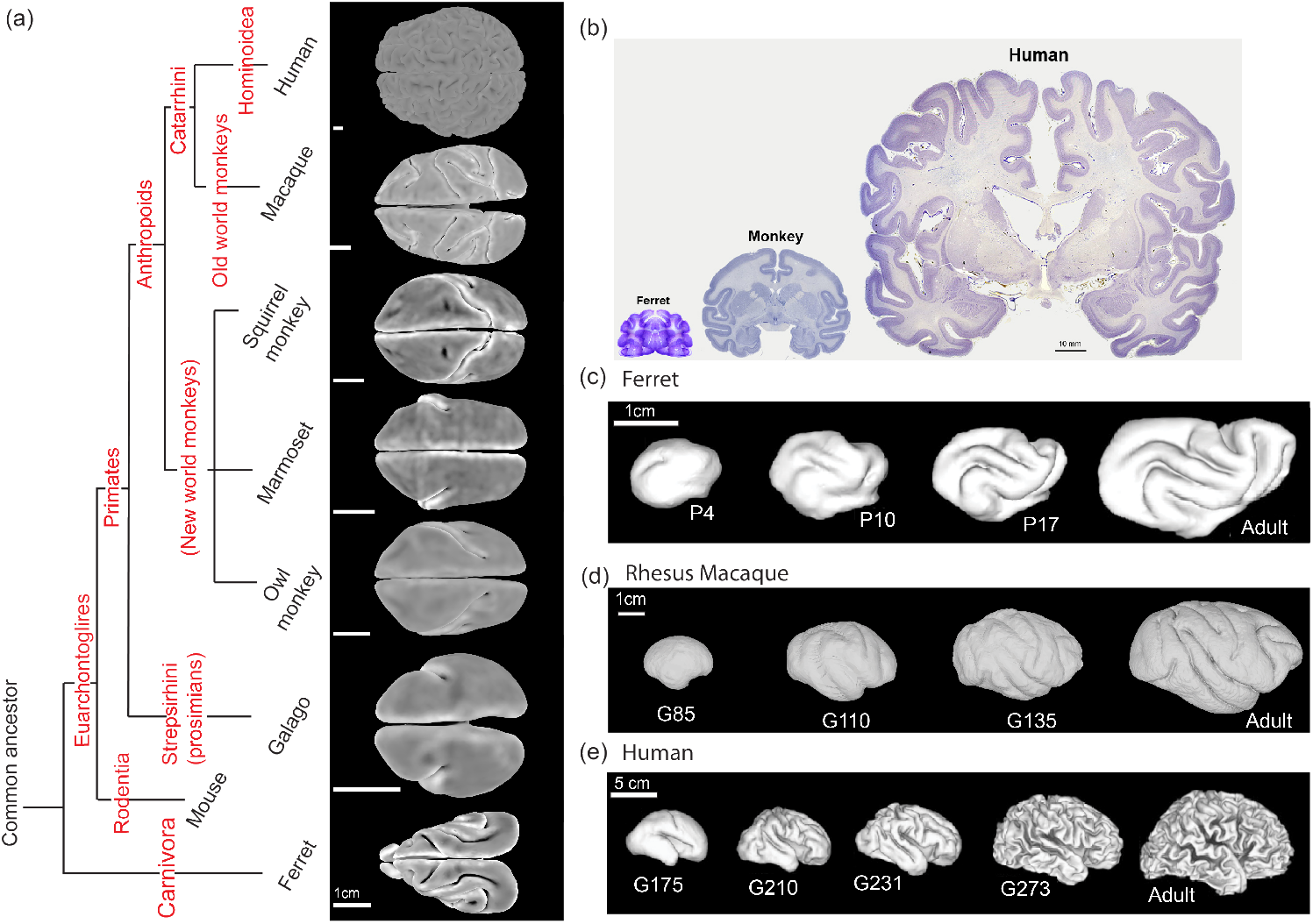
The diversity of the cortical morphologies and developmental processes across species. (a) Phylogenetic relationship of species. Adapted from ***Takahata et al. (2012); Heuer et al. (2019***). Typical real brain surfaces of ferret and primates are presented. Color represents mean curvature. Scale bars: 1 cm (estimated from ***Herculano-Houzel (2009***)). (b) Stained sections of mature brain tissue from ferret, rhesus macaque, and human. Scale bar: 10 mm. Adapted from ***Noctor (2016***). (c-e) 3D reconstruction of cortical surfaces of ferret, macaque, and human brains from fetal to adult. (c) Ferret: postnatal day 4, 10, 17 and adult maturation ***Barnette et al. (2009***). Scale bar: 1cm; (d) Macaque: gestation day 85, 110, 135 ***Liu et al. (2020***), and adult maturation ***Calabrese et al. (2015***). Scale bar: 1cm; (e) Human: gestation day 175 (week 25), 210 (week 30), 231 (week 33), 273 (week 39), and adult maturation ***Barnette et al. (2009***). Scale bar: 5cm.

The development of cortical morphology involves the coordinated and localized expression of many genes that lead to the migration and differentiation of neural stem and progenitor cells ***van der Meer and Kaufmann (2022); Oegema et al. (2020); Del-Valle-Anton and Borrell (2022***). All these biological processes cooperatively generate an expansion of the cortex relative to the underlying white matter and eventually drive cortical folding ***Akula et al. (2023***). While a range of mechanisms have been proposed in the past for the processes leading to folding ***Striedter et al. (2015); Holland et al. (2015); Van Essen (2020***), over the past decade, theoretical and experimental evidence have converged on the primary determinant of folding as a mechanical instability associated with the formation of a localized crease or sulcus ***Toro and Burnod (2005); Hohlfeld and Mahadevan (2012); Tallinen et al. (2013***) driven by differential growth, with iterations and variations that qualitatively explain the brain gyrification ***Tallinen et al. (2014***, 2016). However, combining this mechanistic model with a comparative perspective that aims to quantify the variability of folding patterns across species, or linking it to genetic perturbations that change the relative expansion of the cortex remain open questions. In a companion paper ***Choi et al. (2025***), we address the latter using the ferret as a model organism, while in the current study, we address the former question using a combination of physical experiments with gel swelling, numerical simulations of differential growth and geometric morphometrics to compare brain morphogenesis in the ferret, the macaque, and the human. The species were selected as they have very different brain sizes and folding patterns (Fig. 1(b)) and thus represent different branches in the evolutionary tree, Carnivora, Old World monkeys, and Hominoidea. Furthermore, in all three species, we have access to a time course of the development of the folds, as shown in Fig. 1(c).

### Experiments on swelling gel-brains

To mimic the mechanical basis for brain morphogenesis based on the differential growth of the cortex relative to the white matter, we used the swelling of physical gels that mimic the fetal brain developmental process during post-gestation stages. Previously, this principle has been demonstrated for *Homo sapiens* (human) brain development ***Tallinen et al. (2016***, 2014). To demonstrate that the same principle applies to other species, we constructed physical gels from the fetal brain MRI scans for *Macaca mulatta* (rhesus macaque) and *Mustela furo* (ferret). In brief, a two-layer PDMS gel is constructed from the 3D fetal brain MRI reconstruction and immersed in an organic solvent. Immersion causes solvent imbibition into the surface of the physical gel which swells leading to a compressive strain in the outer layers that causes the surface layer to form convolutions that resemble the folding patterns in the brain cortex layer. Time-lapse images of the gel model to mimic brain folding in *Macaca mulatta* (rhesus macaque) up to G110 are shown in Fig. 2(a), while in Fig. 2(b), we show the initial and final states of swelling to mimic different post-gestation stages corresponding to G85, G110 and G135. A visual inspection of the swollen gels constructed from different post-gestation stages showed qualitatively different folding patterns, indicating the sensitivity of the folds to the initial undulations present on the physical gel surfaces. In Fig. 2(c) and Movie S1, we show the results of similar physical gel experiments to mimic brain folding morphogenesis for *Homo sapiens* ***Tallinen et al. (2016***) and *Mustela furo* ***Choi et al. (2025***); in each case the initial condition was based on 3d fetal brain MRIs and the final state was determined qualitatively using the overall volume of the brain relative to the initial state. No attempt was made to vary the swelling ratio of the surface as a function of location, although it is likely that in the different species this was not a constant. To quantitatively describe the folding patterns, the swollen gel surfaces were then scanned and reconstructed using X-ray Computed Tomography (Methods). Movie S2–S4 showed the fetal brain MRI scans and the reconstructed 3D swollen gel surfaces in juxtaposition for *H. sapiens, M. furo*, and *M. mulatta*. This paves the way for a quantitative comparison of the results of the physical experiments with those derived from a mechanical theory for brain morphogenesis and those from scans of macaque, ferret and human brains.

**Figure 2.**
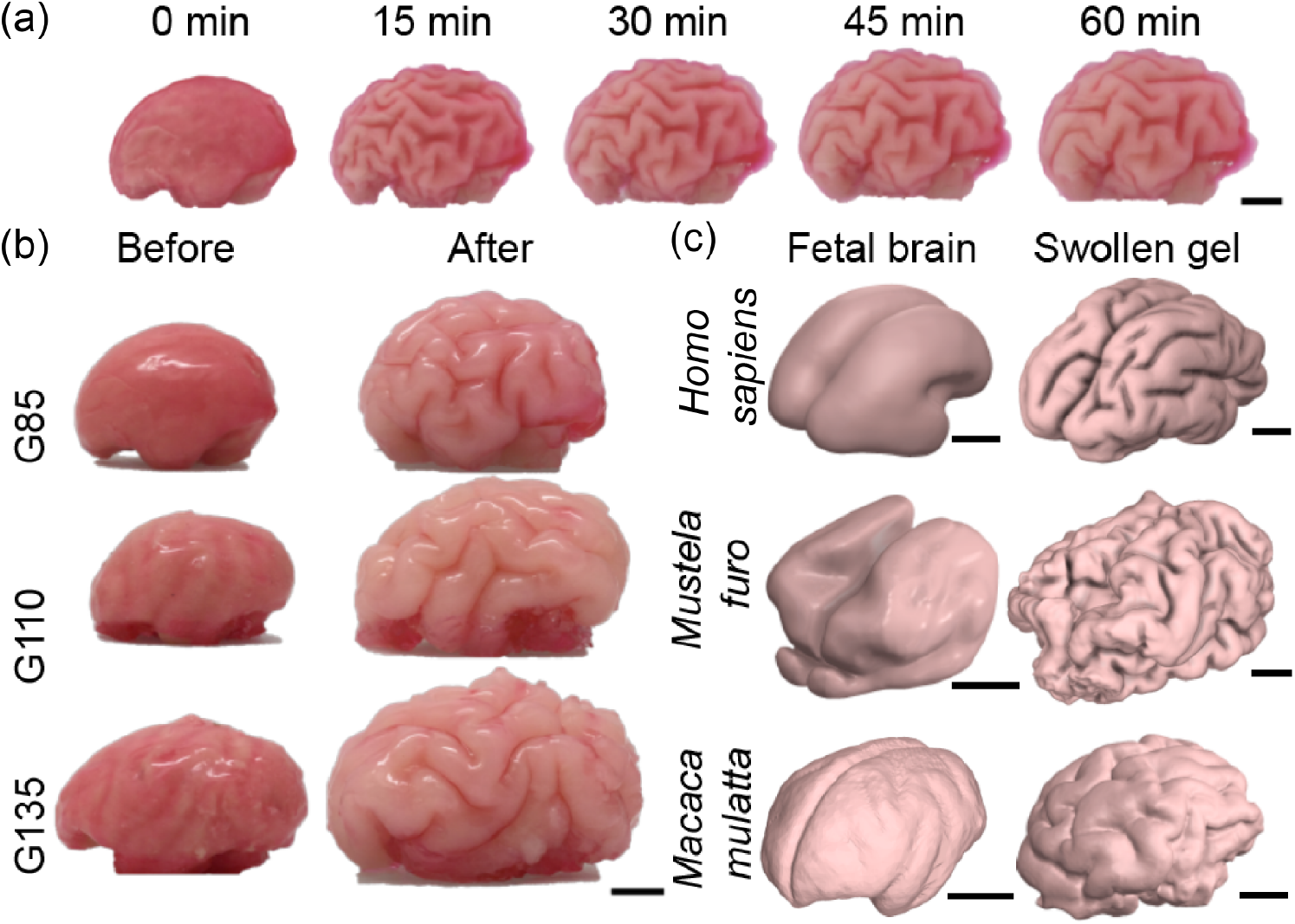
Physical gel model that recapitulates the growth-driven morphogenesis mechanism across phylogeny and developmental stages. (a) A time-lapse of the physical gel brain mimicking macaque brain development starting from G110. (b) Left views of three physical gels mimicking macaque post gestation day 85, day 110, and day 135 before and after hexane swelling. Scale bar: 1 cm. (c) Comparison of fetal/newborn brain scans and the reconstructed surfaces of swollen physical gels for various species. Scale bars: 1 cm.

### Simulations of growing brains

To test the capability of the mechanical model for brain morphogenesis based on differential growth ***Tallinen et al. (2014); Budday et al. (2014); Tallinen et al. (2016***) to explain patterns across species, we perform numerical simulations of the developing brains of ferret, macaque, and human modeled as soft tissues. Here we only consider the simplest homogeneous growth profile which is sufficient to capture the folding formation across different species; regional growth of the cortical layer based on real data from tracking the surface expansion of fetal brains ***Garcia et al. (2018); Alenyà et al. (2022); Weickenmeier (2023***) is a more sophisticated alternative that we do not adopt for reasons of simplicity.

The initial brain models are reconstructed from 3D fetal brain MRI (*Methods*), and assumed to be composed of gray and white matter layers which are considered as hyperelastic materials with differential tangential growth ratio *g*. A multiplicative decomposition of deformation gradient gives F = A·G with A the elastic part and 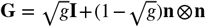 the growth tensor. We adopt a modestly compressible neo-Hookean material with strain energy density

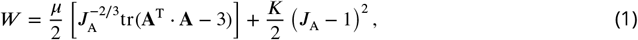

where F is the deformation gradient, *J*_A_ = det A, *μ* is the initial shear modulus, and *K* is the bulk modulus. Considering modestly compressible material, we assume *K* = 5*μ*.

We solve the final shapes of the growing tissues using a custom finite element method ***Tallinen et al. (2016***). All the initial geometries of smooth fetal brains are obtained from open data sources and through collaboration (*Methods*, 3D model reconstruction), and the growth ratio distribution is assumed as a function of the initial location, including the distance to the cortical surface and the presumably non-growing regions. Other parameters, such as the thickness ratio and modulus ratio of gray and white matter, and the temporal changes of growth ratios are assumed (*Methods*, numerical simulations, Table 2). The brain models are discretized to tetrahedrons by Netgen ***Schöberl (1997***). An explicit algorithm is adopted to minimize the total strain energy of the deforming tissues. We adopt a step-wise simulation strategy where the initial geometry of each presumed stress-free state is obtained from real MRI data of earlier-stage fetal brains, instead of using a continuous model where only the initial smooth brain geometry is input ***Tallinen et al. (2016***). The simulated developmental processes of fetal brains are presented in Fig. 3 and Movie S1.

**Figure 3.**
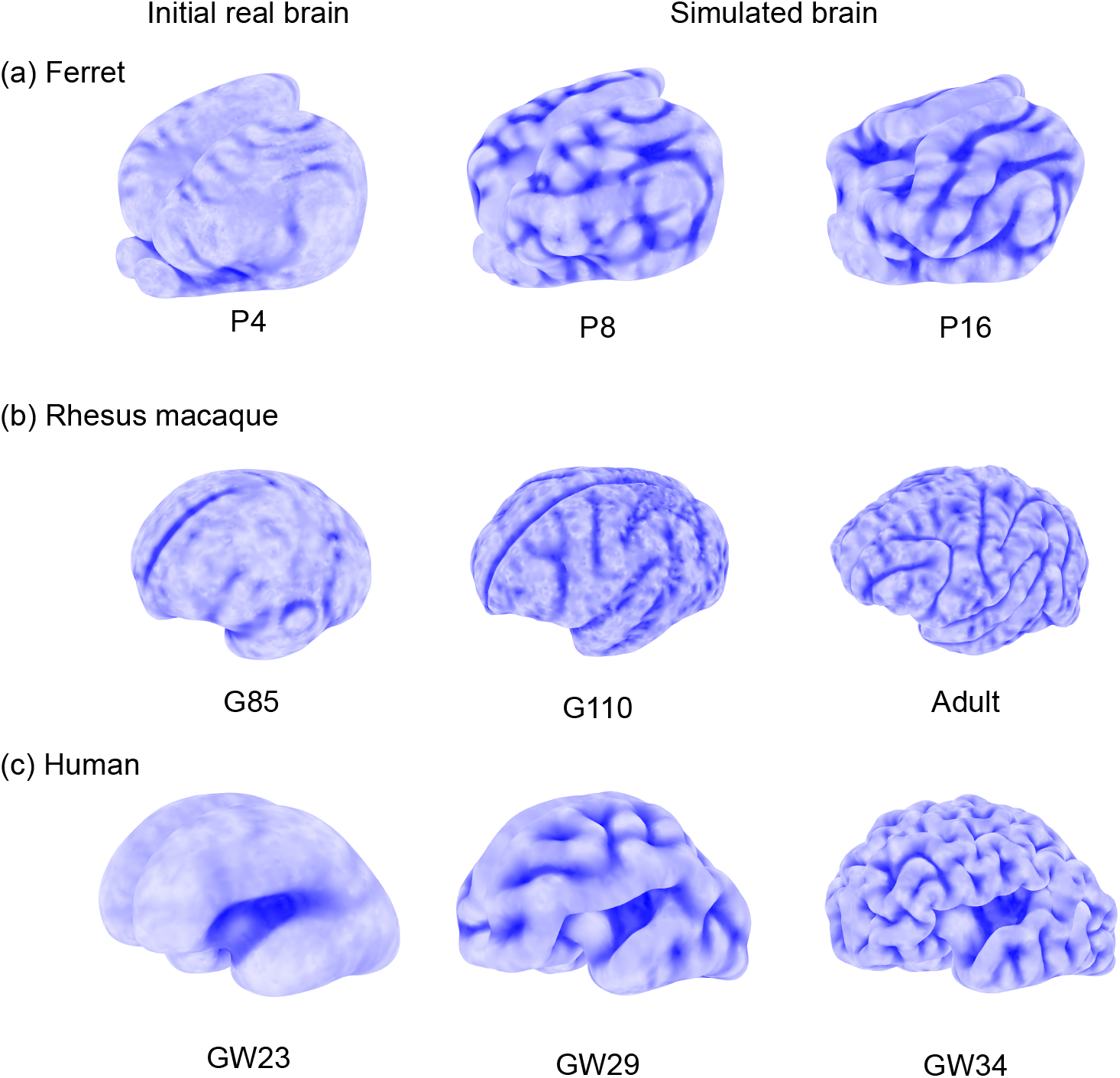
Simulations of growing brains of. (a) ferret, (b) rhesus macaque, and (c) human. Starting from smooth fetal/newborn brains, simulations show different gyrification patterns across species. The brains are modeled as soft elastic solids with tangential growth in the gray matter (see *Simulations of growing brains* for details). Initial 3D geometries are taken from the reconstruction of MRI (see *Methods*, 3D model reconstruction). Mechanical parameters of growth ratio and cortical thickness are provided in Table 2. Color from dark to light blue represents shape index (as defined in Eq. (2)) from −1 to 1.

### Morphometric analysis

To verify whether our simulation methods and physical gel models are sufficient to capture the developmental process of fetal brains and reproduce cortical patterns in adult brains, we compare the cortical surfaces of real, simulated, and gel brains across different species. Fig. 4(a) presents the hemispherical cortical surfaces of the real, simulated, and gel brains (denoted as S_1_, S_2_, and S_3_, respectively). Left and right symmetry and the comparison of whole brains are presented in Fig. S3 and S4, Movie S2–S4. Major sulci are extracted and highlighted by hand for further alignment. To analyze the shape differences, we then adapted the parameterization methods in ***Choi and Lui (2015); Choi et al. (2015***, 2020) to map the brain surfaces onto a common disk shape with the major sulci aligned using landmark-matching disk quasi-conformal parameterizations. Denote the three disk parameterization results as *D* _1_, *D* _2_, *D* _3_, as shown in Fig. 4(b). Multiple quantitative measures such as surface area, cortical thickness, curvature, and sulcal depth can be adopted to analyze curved surfaces and to compare different surfaces. Here, we use shape index (SI) and rescaled mean curvature 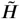 to describe the similarity among real, simulated, and gel surfaces. Shape index is a dimensionless and scale-independent surface measure ***Koenderink and van Doorn (1992***). It can be calculated from the mean and Gaussian curvatures by

**Figure 4.**
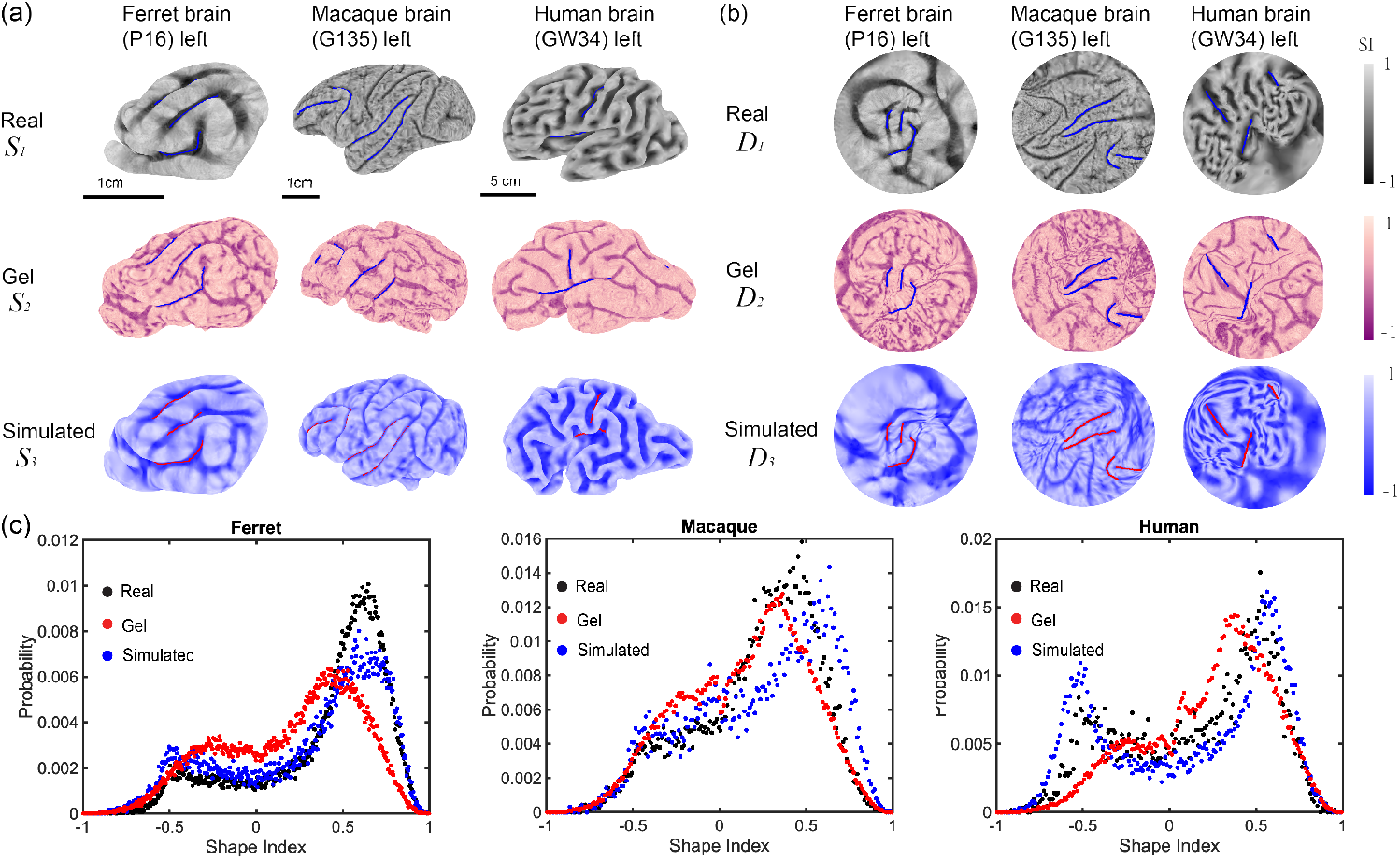
Comparison among real (*S* _1_), simulated (*S*_2_), and gel brains (*S*_3_) of ferret, rhesus macaque, and human via morphometric analysis. (a) 3D cortical surfaces of *in vivo, in vitro* and *in silico* models. Left brain surfaces are provided here. The symmetry of the left and right halves of the brain surfaces is discussed in Figs. S3 and S4, Movie S2–S4. Three or four major folds of each brain model are highlighted and served as landmarks. The occipital pole region of macaque brains remains smooth in real and simulated brains. (b) The quasi-conformal disk mapping with landmark matching of cortical surfaces on disk (see Sec *Morphometric analysis* for details). Blue or red curves represent corresponding landmarks. Color represents shape index (SI, as defined in Eq. (2)). Similarity indices of each simulated and gel brain surfaces are presented in Table 1. (c) Histogram of shape index of ferret, macaque, and human. Black, red, and blue dots represent the probability of shape index of real, gel, and simulated surfaces, respectively.

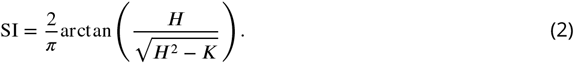

**Table 1.**
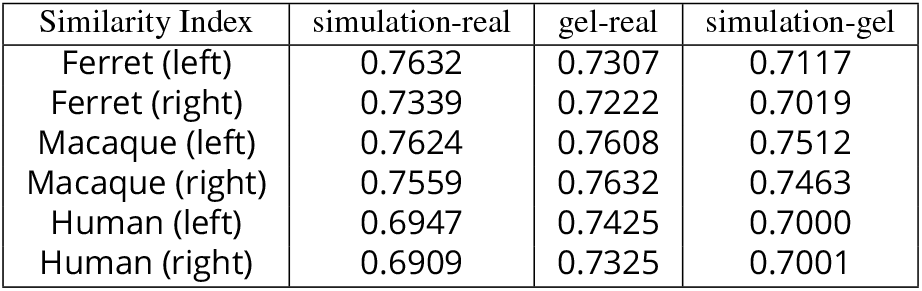
Similarity index evaluated by comparing the shape index of simulated brains (S), swollen gel brain simulacrums (G) and real brain surfaces (R), calculated with vector p-norm *p* = 2, as described in Eq. (4).

**Table 2.**
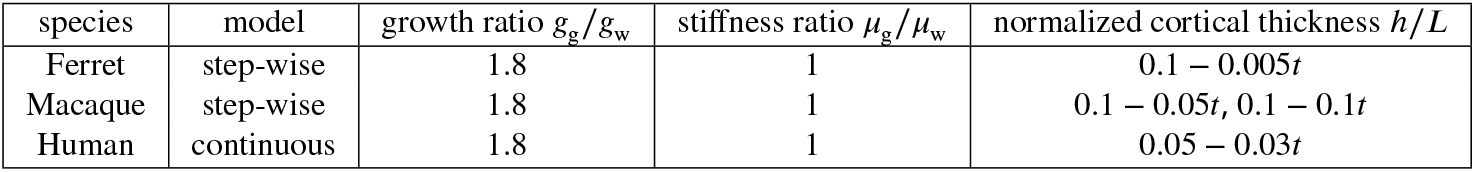
Parameters for numerical simulations.

| species | model | growth ratio $g_g/g_w$ | stiffness ratio $\mu_g/\mu_w$ | normalized cortical thickness $h/L$ |
| --- | --- | --- | --- | --- |
| Ferret | step-wise | 1.8 | 1 | $0.1 - 0.005t$ |
| Macaque | step-wise | 1.8 | 1 | $0.1 - 0.05t, 0.1 - 0.1t$ |
| Human | continuous | 1.8 | 1 | $0.05 - 0.03t$ |

Shape index ranging within [−1, 1] defines a continuous shape change from convex, saddle, to concave shapes. Fig. S2 shows nine categories of typical curved surfaces. For brain surfaces, we can classify sulcal pits (−1 < SI < −0.5), sulcal saddles (−0.5 < SI < 0), saddles(SI = 0), gyral saddles (0 < SI < 0.5) and gyral nodes (0.5 < SI < 1). When the shape index equals −1 or 1, it represents a defect of the curvature tensor with two eigenvalues identical, as shown in Fig. S3. The probability of shape index distribution exhibits two peaks, as shown in Fig. 4(c), corresponding to ridge and rut shapes (SI=±0.5), where the ridge shape (SI= 0.5) is dominating. In contrast, the rescaled mean curvature histogram exhibits a unique peak around 0.2 (Fig. S3). The two-peak and unique-peak distributions of adult human brain surfaces have also been presented in previous research ***Demirci and Holland (2022); Hu et al. (2013***). To quantify the similarities between every two brain surfaces, we evaluate the distribution of *I*(*υ*) differences on the common disk domain at each vertex *υ*. Here I(*υ*) represents either the surface shape index SI(*υ*) or the rescaled mean curvature 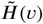 defined as

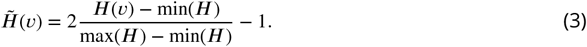

Thus the similarity *s* of the distributions 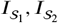 of the two surfaces *S*_1_, *S*_2_ is then evaluated via

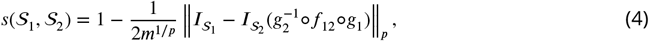

where *m* is the total number of vertices, *g*_1_ : *S*_1_ → *D*_1_ and *g*_2_ : *S*_2_ → *D*_2_ are the initial disk conformal parameterizations, *f*_12_ is the landmark-aligned quasi-conformal map between *D* and *D*_2_, and ∥·∥ is the vector p-norm:

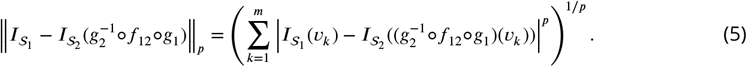

Note that since 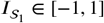 and 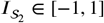, we have 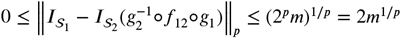.

Therefore, we always have 0 ≤ *s* (*S*_1_, *S*_2_) ≤ 1.

We calculate the similarity indices with different p-norm: *p* = 1, *p* = 2, and *p* = ∞ of both rescaled mean curvature and shape index. The results are given in Table 1 and Table S2.

## Discussion

In this study, we have explored the mechanisms underlying brain morphogenesis for a few different mammalian species. Using fetal and adult brain MRIs for ferrets, macacques and humans, we carried out physical experiments using swelling gels, combined with a mathematical framework to model the differential growth of the cortex that leads to iterations and variations of a mechanical (sulcification) instability that allowed us to recapitulate the basic morphological development of folds. We then deployed a range of morphometric tools to compare the results of our physical experiments and simulations with 3d scans of real brains, and show that our approaches are qualitatively and quantitatively consistent with experimental observations of brain morphologies.

All together, our study shows that differential growth between the gray matter cortex and white matter bulk provides a minimal physical model to explain the variations in the cortical folding patterns seen in multiple species. More specifically, we see that the overall morphologies are controlled by the relative size of the brain (compared to the cortex), as well as the scaled surface expansion rate, both of which can and do have multiple genetic antecedents ***Bae et al. (2014); Shinmyo et al. (2017); Qi et al. (2023); Lai et al. (2003); Barresi et al. (2024***) (see SI, Table S1).

Our results point to some open questions. First, the relationship between physical processes that shape organs and the molecular and cellular processes underlying growth has been the subject of many recent studies, e.g. in the context of gut development ***Gill et al. (2024a***,b), and it would seem natural to expect similar relationships in brain development. There is a growing literature linking genes with brain malformation and pathologies. For example, GPR56 ***Bae et al. (2014***) and Cdk5 ***Shinmyo et al. (2017***) can affect progenitors and neurons in migration, SP0535 ***Qi et al. (2023***) can affect neural proliferation, and foxpp2 can affect neural differentiation ***Lai et al. (2003); Barresi et al. (2024***), all of which change the cortical expansion rate and thickness, consequently leading to brain malformation and pathologies, as listed in Table S1. While a direct relation between gene expression levels and the effective tangential growth rate G and cortical thickness has only been partially resolved, as for example in our companion study on the folding and misfolding of the ferret brain ***Choi et al. (2025***), further studies are needed to address how genetic programs drive cell proliferation, migration, size and shape change that ultimately lead to different cortical morphologies. Second, although our focus has been exclusively on the morphology of the brain, recent studies ***Pang et al. (2023); Schwartz et al. (2023***) are suggestive of a link between cortical geometry and function from both developmental and evolutionary perspectives, and suggest natural questions for future study. Third, despite prescribing a simple spatially homogeneous form for the cortical expansion for fetal brain surfaces of all three species studied, we were able to capture the essential features of the folds and variations therein. The effect of spatio-temporally varying inhomogeneous growth needs to be further investigated by incorporating regional growth of the gray and white matter not only in human brains ***Garcia et al. (2018); Weickenmeier (2023***) based on public datasets ***Namburete et al. (2023***), but also in other species to investigate folding differences across species, inter-individual variability and finally regional differences in folding. More accurate and specific work is expected to focus on these directions. Finally, our physical and computational models along with our morphometric approaches are a promising avenue to pursue in the context of the inverse growth problem, i.e., postulate the fetal brain morphologies from the adult brains. This may one day soon allow us to reconstruct the adult brain geometries from fossil endocasts ***de Sousa et al. (2023***), and eventually provide insights into how a few mutations might have triggered the rapid expansion of the cortex across evolutionary time and led to the highly convoluted human brain that is able to ponder the question of how it might have folded itself.

## Methods

### 3D model reconstruction

#### Pre-processing

We used a publicly available database for all our 3d reconstructions: fetal macaque brain surfaces are obtained from ***Liu et al. (2020***); newborn ferret brain surfaces are obtained from project FIIND ***Toro et al. (2018***); and fetal human brain surfaces are obtained from ***Tallinen et al. (2016***). These 2D manifolds of brain surfaces were first normalized by their characteristic lengths 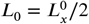 and discretized to triangle meshes by Meshlab ***Cignoni et al. (2008***) and then converted to 3D models and discretized to tetrahedral elements in Netgen https://ngsolve.org/.

#### Post-processing

All the numerically calculated brain surfaces and scanned gel brain surfaces were extracted as 3D triangle meshes. These intact cortical surfaces were then dissected to left and right semi-brains and normalized by half of their longitudinal lengths *L* = *L*_*x*_ /2. These surfaces were then checked and fixed to a simply connected open surface to satisfy the requirements for open disk conformal mapping ***Choi and Lui (2015***).

### Experimental protocol for gel experiments

The physical gel was constructed following a previous publication ***Tallinen et al. (2016***). In brief, a negative rubber mold was generated with Ecoflex 00-30 from a 3D-printed fetal brain plastic model. The core gel was then generated with SYLGARD 184 at a 1:45 crosslinker: base ratio. Three layers of PDMS gel at a 1:35 crosslinker:base ratio were surface-coated onto the core layer to mimic the cortex layer. Pigments were added to the PDMS mixture for bright-field visualization. To mimic the cortex folding process, the physical gel was immersed in n-hexane. The time-lapse videos were taken with an iPod Touch 7th Gen. To reconstruct the swollen gel surface and analyze the folding patterns, swollen gels were imaged with a Nikon X-Tek HMXST X-ray CT machine. The voxel resolution for all scans was 100 μm. To minimize solvent evaporation during the 30-minute scan, cotton soaked in the solvent was placed inside the container. The container’s thin acrylic walls allowed for clear X-ray transmission. The container’s thin acrylic walls allowed for clear X-ray transmission. To test the reversibility of the folding pattern formation, the physical gel models were allowed to dry in a laminar flow hood overnight before being immersed in hexane again.

### 3D reconstruction of swollen gel surface from X-ray CT

The z-stack images obtained from the X-ray CT machine were segmented by a machine-learningbased segmentation toolbox, Labkit, via ImageJ ***Schneider et al. (2012***). A classification was created for each swollen gel. Then a 3D surface of the segmented gel image was generated by ImageJ 3D viewer. Further post-processing was conducted in SOLIDWORKS and MeshLab ***Cignoni et al. (2008***), including reversing facial normal direction, re-meshing, and surface-preserving Poisson smoothening.

### Numerical Simulations

The simulation geometries of ferret, macaque, and human are based on MRI of smooth fetal or newborn brains. For the ferret, we take the P0, P4, P8, and P16 postnatal brains as initial shapes of the step-wise growing model; for the macaque, we take G80 and G110 fetal brains as initial shapes of the step-wise growing model. To focus on fold formation, we did not consider the lack of patterning in the relatively smooth regions, such as the Occipital Pole of the macaque; for the human, we take the GW22 fetal brain as the initial shape of the continuous growing model. Small perturbations of the initial geometry typically affect only the minor folds, while the main features of the major folds, such as their orientation, width, and depth, are well conserved across individuals ***Bohi et al. (2019); Wang et al. (2021***). For simplicity, we do not perturb the fetal brain geometry obtained from datasets. Both gray and white matter are considered as neo-Hookean hyperelastic material with the shear modulus distribution

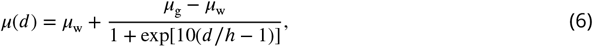

where *d* is the distance from an arbitrary material point inside the brain to the cortical surface, and *h* is the approximated cortical thickness assumed to decrease with growth process time *t* ***Tallinen et al. (2016***). The subscripts ‘g’ and ‘w’ represent gray and white matter, respectively. The tangential growth ratio *g* has a similar spatial distribution

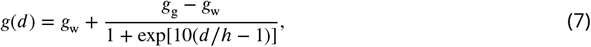

where *g*_w_ and *g*_g_ represent the growth ratio at cortical surface and at innermost white matter. The parameters used in simulations are listed in Table 2.

## Data and code availability

Reconstructed 3D surface models of fetal and adult brains of macaque and human are available on GitHub at https://github.com/YinSifan0204/Comparative-brain-morphlogies. Requests for the ferret brain data should be made to the team of Roberto Toro and Katja Heuer. All other data are included in the article and/or supplementary material.

## Author contributions

L.M. and R.T. conceived the project; S.Y., C.L., G.P.T.C., and L.M. designed research; S.Y. collected data and performed numerical simulations; C.L. and Y.J. performed gel experiments; S.Y. and G.P.T.C contributed morphometric analysis; K.H. and R.T. provided developmental ferret and adult primate brain data; S.Y., G.P.T.C., and L.M. wrote the paper with input from all authors; all authors edited the paper; L.M. supervised the project.

## Acknowledgements

We acknowledge partial financial support from the NSF-ANR grant 2204058 (L.M., K.H., R.T.), the Simons Foundation and the Henri Seydoux Fund (L.M.), the CUHK Faculty of Science Direct Grant for Research #4053650 (G.P.T.C.), the project NeuroWebLab (ANR-19-DATA-0025, K.H., R.T.), DMOBE (ANR-21-CE45-0016), the European Union’s Horizon 2020 research and Marie Skłodowska-Curie grant agreement 101033485 (K.H.).

## Supplementary material

**This PDF file includes**

Supplementary Text

Figs. S1 to S5

Movies S1 to S4 descriptions

## Supplementary Text

### Malformations of human cortical development

In the main text, we have shown examples of normal cortical development and the resulting folding processes by following fetal and adult animals across a few different species. Defects in 1neuronal migration disorders may lead to a group of rare brain malformations, the most severe forms being lissencephaly and polymicrogyria, as shown in Fig. S1(a) (*15*). Lissencephaly, or agyria-pachygyria, is characterized by a simplified convolutional pattern where a few broad gyri are separated by rudimentary primary fissures and sulci. Lissencephaly is accompanied by a very thick cortical gray matter layer (Fig. S1(a), middle) which has been confirmed as a critical cause in this malformation (*3,18*). Polymicrogyria, on the contrary, is an overly convoluted cortex with a reduced cortical thickness and an increasing number of secondary folds (Fig. S1(a), right). Various malformations of cortical development are congenital and genetically heterogeneous diseases which mutations or deletions of genes have been identified. For example, a regional deletion mutation in a regulatory element of GP56 can selectively disrupt the human cortex surrounding the Sylvian fissure bilaterally including “Broca’s area”. Fig. S1(b) shows the MRI of polymicrogyria of a noncoding mutation in the GPR56 gene (*1*). This abnormally thin cortex is folded giving a paradoxical but characteristic thickened appearance. Many other genes have also been found associated with brain malformation and pathologies, such as foxp2 (*2,11*), SP0535 (*16*) and Cdk5 (*17*). These genes affect progenitors and neurons in migration, proliferation, and differentiation which change the cortical expansion rate and thickness, consequently leading to brain malformation and pathologies, as listed in Table S1.

### Statistical characteristics of brain folding

To characterize the form of the convolutions on the surface of the brain, we use a dimensionless and scale independent surface measure, the shape index (SI), to quantify 3D cortical morphologies across species. The definition of shape index is given as (*9*)

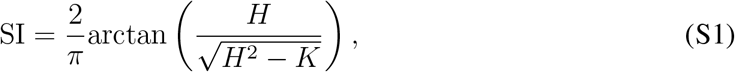

or equivalently,

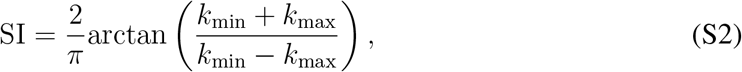

where *k*_max_ and *k*_min_ are the maximum and minimum curvatures, respectively,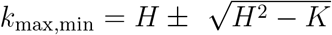. The shape index is a continuous number representing the local shape of a surface. Fig.S2 illustrates a shape index scale divided into nine categories: spherical cup, trough rut, saddle rut, saddle, saddle ridge, ridge, dome and spherical cap, with the color representing the shape index value ranging from −1 to 1.

In the main text, we have shown the shape index distribution of left semi-sphere brains.

Here, we present the results of both left and right semi-spheres of ferret, macaque and human brains in Fig. S3 (top two rows) and Fig. S4 with comparison among the real (black dots), gel (red dots), and simulated (blue dots) surfaces. The probability of shape index distribution exhibits two peaks, corresponding to ridge and rut shapes (SI=±0.5), where the ridge shape (SI=0.5) dominates. In contrast, the rescaled mean curvature histogram exhibits a unique peak around 0.2 (Fig. S3, the 3rd and 4th rows). These histograms illustrate that shape index and mean curvature exhibit qualitatively different distributions and both of these should be taken into account when describing cortical surface characteristics, consistent with earlier studies (*7, 8*).

### Morphometric analysis

For the morphometric analysis, we developed a landmark-matching disk quasi-conformal parameterization method by extending and combining the methods in (*5, 6*) and applied it to map the real, simulated, and gel brain surfaces onto the unit disk with major sulci aligned.

More specifically, we denote the real, simulated, and gel brain surfaces as *S*_1_, *S*_2_, *S*_3_ respectively, and note that *S*_1_, *S*_2_, *S*_3_ are all simply connected open surfaces and hence are topologically equivalent to the unit disk. We then started by applying the disk conformal parameterization method in (*6*) to map *S*_1_, *S*_2_, *S*_3_ onto the unit disk, followed by the Möbus area correction scheme in (*4*) to further reduce the area distortion of the mappings. Denote the three disk parameterization results as *D*_1_, *D*_2_, *D*_3_.

Next, we followed the idea in (*5*) to compute landmark-aligned quasi-conformal maps between *D*_1_, *D*_2_, *D*_3_. More explicitly, given certain major sulci identified on each of the three brain surfaces *S*_1_, *S*_2_, *S*_3_, we computed two landmark-matching quasi-conformal maps *f*_13_ : *D*_1_ → *D*_3_ and *f*_23_ : *D*_2_ → *D*_3_ that deformed *D*_1_ and *D*_2_ to align the sulci on them with the corresponding sulci positions on *D*_3_. Consequently, we can visualize and compare the folding patterns of the three brain surfaces S_1_, S_2_, S_3_ by using the three disk parameterizations *f*_13_(*D*_1_), *f*_23_(*D*_2_), *D*_3_.

### Diversity and evolution of cerebral folding in primates

For a surface-based (superficial) comparison of the brain morphologies among different primates from an evolutionary perspective, we take advantage of the fact that biologists have provided plenty of anatomical data on brain geometry, although a systematic study on how these cerebral folding patterns vary across different sizes, initial shapes, and foldedness for human and other primate species is still lacking. Among primates, brains vary enormously from roughly the size of a grape to the size of a grapefruit, and from nearly smooth to dramatically folded; of these, human brains are amongst the most folded, and the largest (relative to body size). These variations in size and form make comparative neuroanatomy a rich resource for investigating common trends that transcend differences between species. We examined 10 primate species in order to cover a wide range of sizes and forms, as shown in Fig. S5. Using our developed morphometric method, more investigations on the scaling law of their cortical thickness relative to the surface geometry, folding with respect to size (total surface area) and geometry (i.e. curvature, shape, and sulcal depth), and foldedness (gyrification) would be expected.

## Movie Captions

**Movie S1**

Part 1: the swelling processes of physical gel brains of ferret, macaque, and human brains; Part 2: the simulations mimicking the developmental processes of fetal brains of ferret, macaque, and human from smooth surface to the convoluted pattern.

**Movie S2**

Comparison of the real (gray), gel (pink) and simulated (blue) ferret brains. Color represents shape index.

**Movie S3**

Comparison of the real (gray), gel (pink) and simulated (blue) macaque brains. Color represents shape index.

**Movie S4**

Comparison of the real (gray), gel (pink) and simulated (blue) human brains. Color represents shape index.

---

**Fig. S1.**
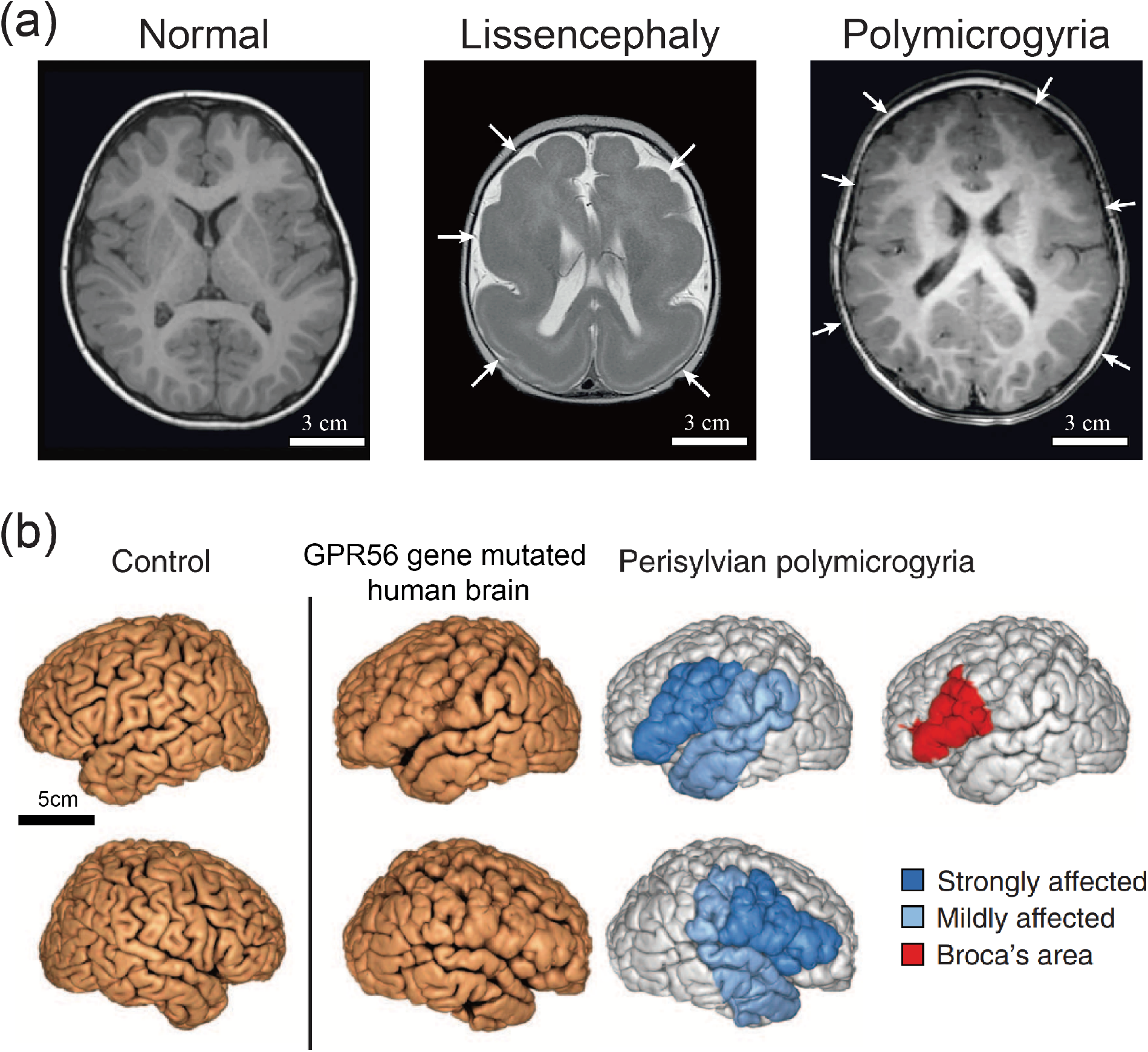
(a) MRI scans showing common malformations of cortical development of human brains. Adapted from (*15*) with permission. Left: normal brain. Middle: lissencephaly spectrum with agyria–severe pachygyria (arrows). Right: bilateral frontoparietal polymicrogyria with abnormally small gyri and shallow sulci (arrows). Scale bars: 3 cm (estimated from (*14*)). (b) A noncoding mutation in the GPR56 gene disrupts perisylvian gyri. MRI shows polymicrogyria in the perisylvian area, resulting in a characteristic, thickened appearance. Adapted from (*1*) with permission.

**Fig. S2.**
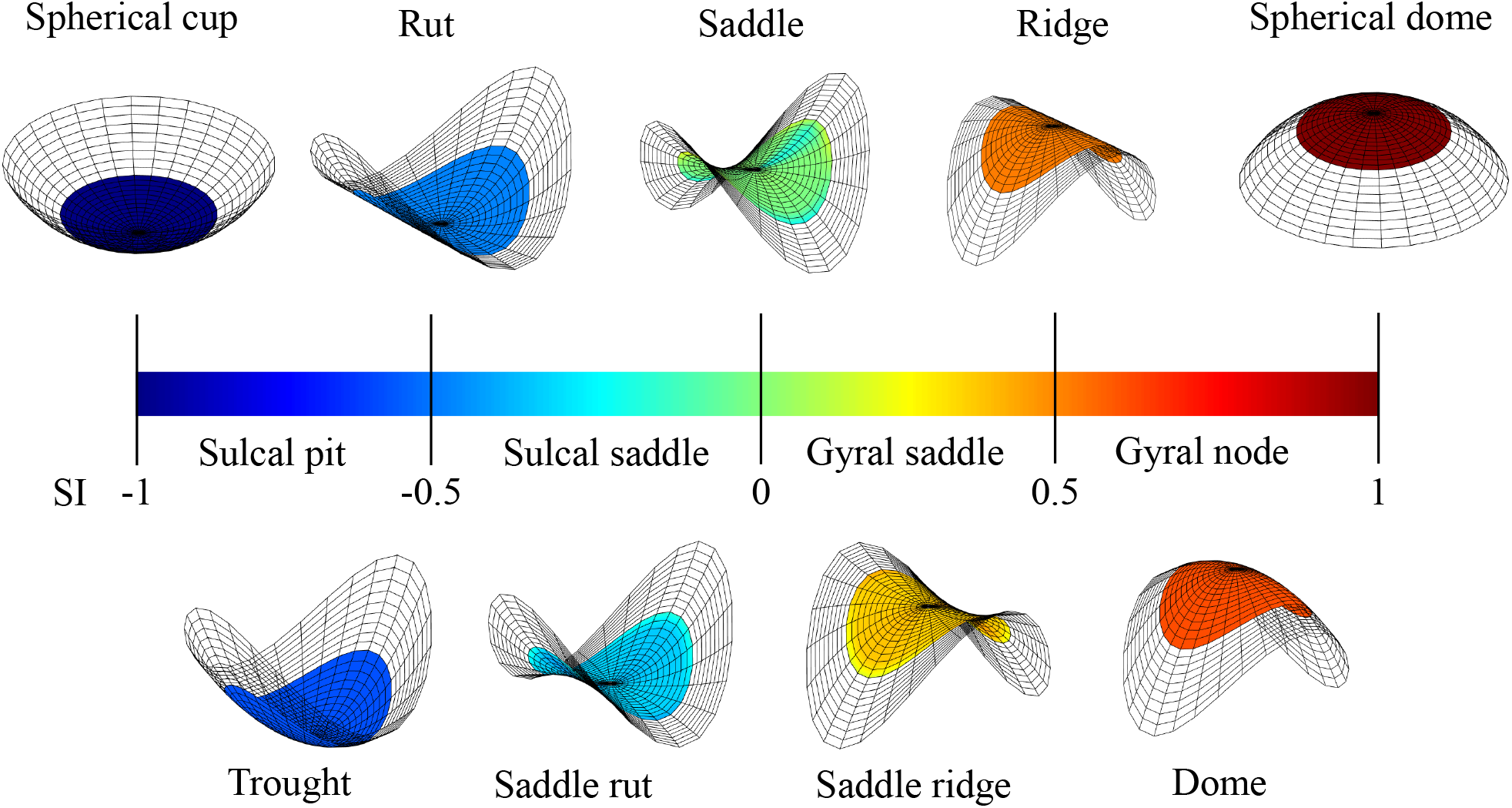
Illustration of shape index scale divided into nine categories: spherical cup, trough rut, saddle rut, saddle, saddle ridge, ridge, dome and spherical cap. The insets are schematics of local curved surfaces. All outward normals pointing upwards.

**Fig. S3.**
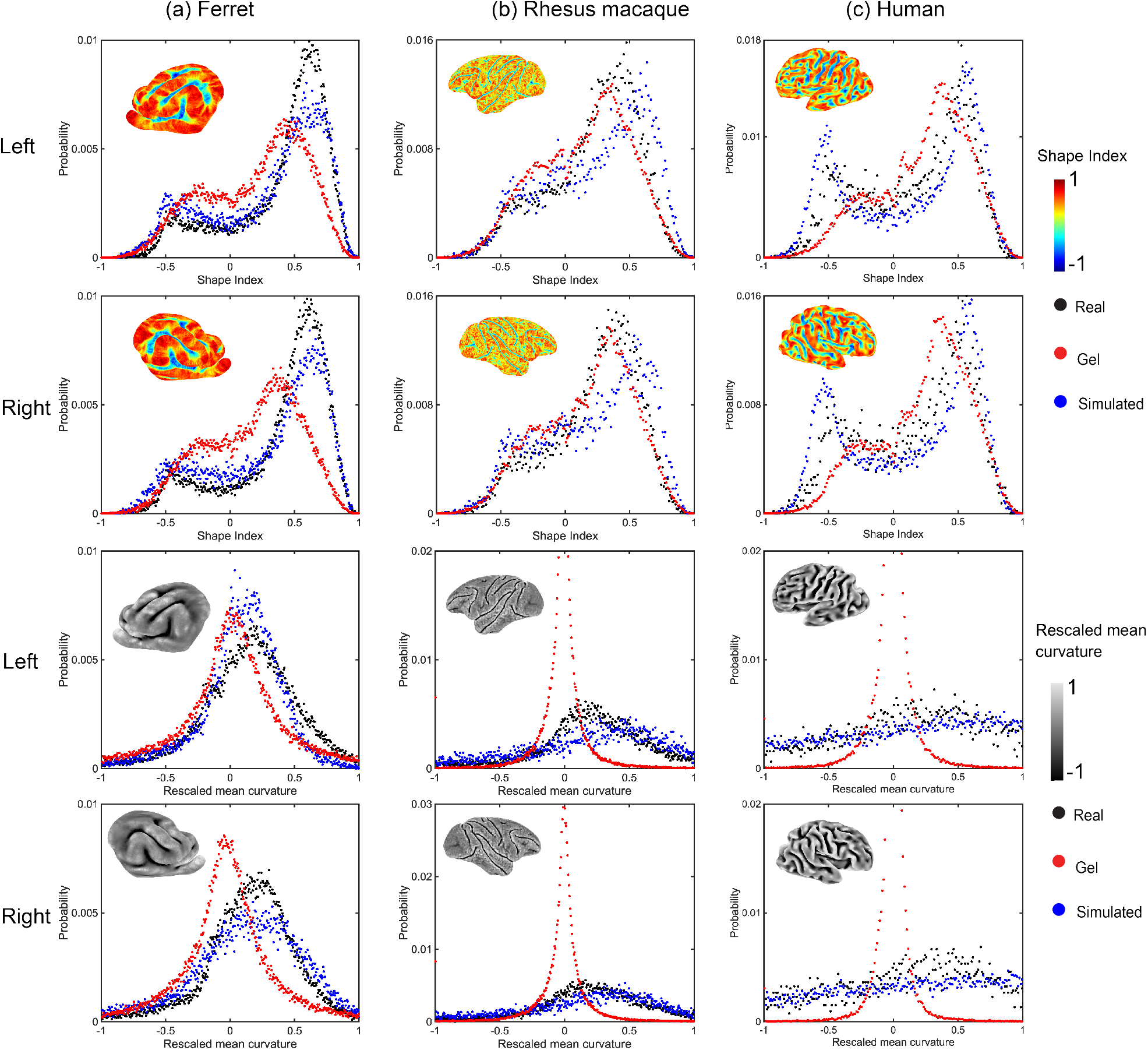
The histogram of shape index SI (top two rows) and rescaled mean curvature 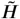 (bottom two rows) of adult cortical surfaces of ferret, macaque and human. Insets are real brain surfaces. Colors represent shape index SI (Eq. (2) in the main text) or rescaled mean curvature 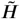 (Eq. (3) in the main text).

**Fig. S4.**
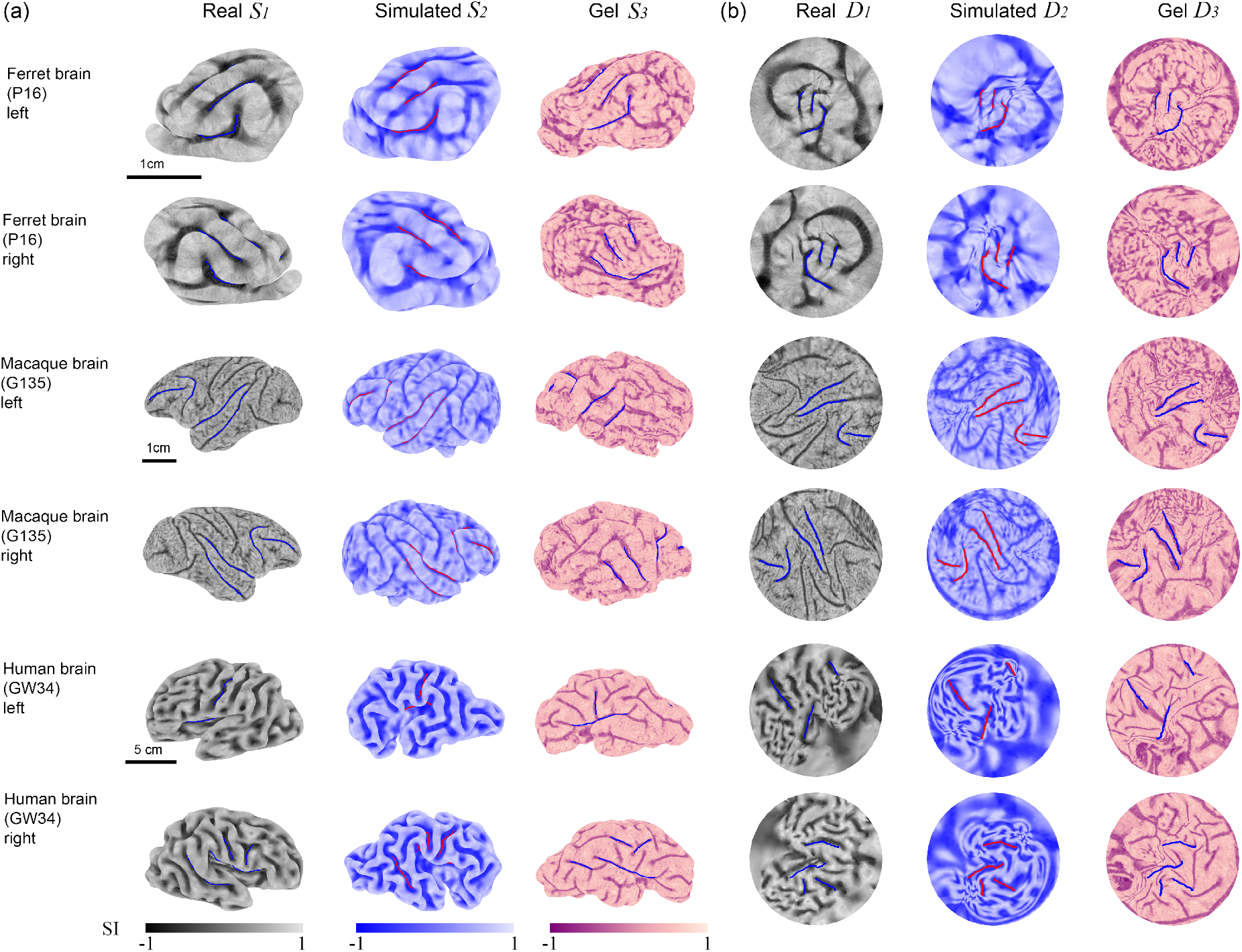
Comparison among real (_1_), simulated (_2_), and gel brains (_3_) of ferret, rhesus macaque, and human via morphometric analysis. (a) 3D cortical surfaces of *in vivo, in silico*, and *in vitro* models. Both left and right cortical surfaces are provided to present the left-right symmetry. (b) The quasi-conformal disk mapping with landmark matching of cortical surfaces on disk. Blue or red curves represent corresponding landmarks. Color represents shape index (SI). Similarity indices of each simulated and gel brain surfaces are presented in Table S2.

**Fig. S5.**
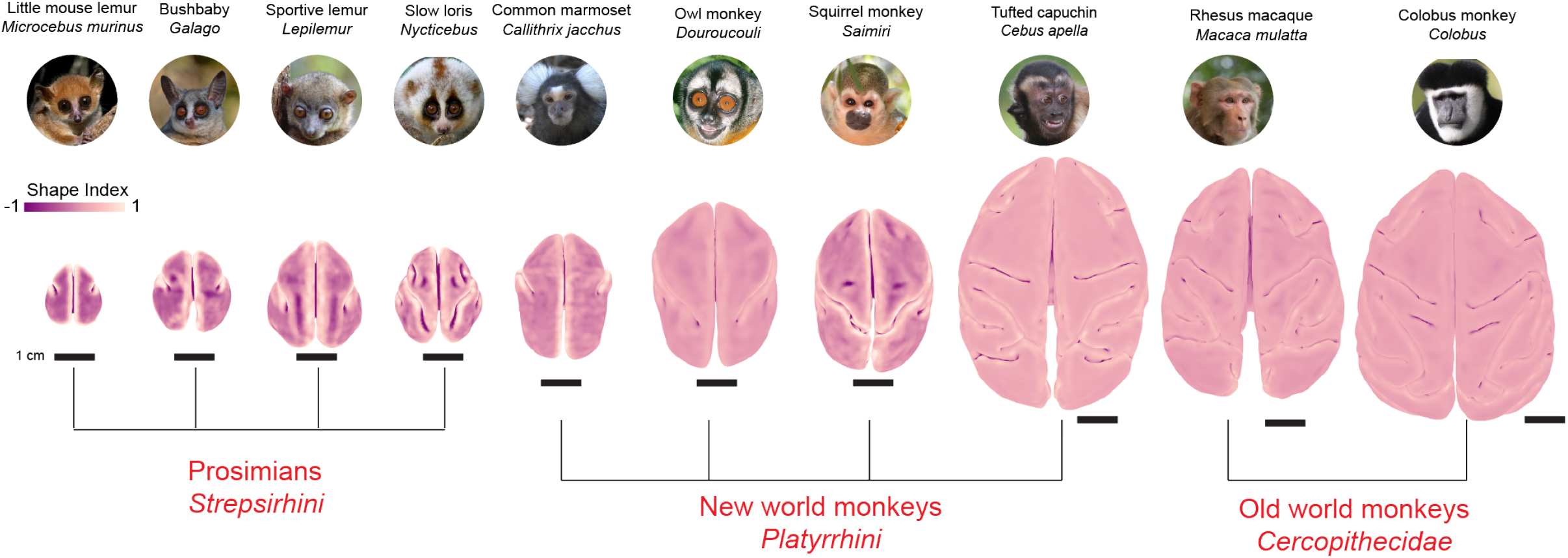
Comparison across ten primate species. Each species is listed by its common name and scientific name, and accompanied by a picture. Scale bar: 1cm. Color represents the shape index. Pictures are taken from Wikipedia.

**Table S1.** Gene-related brain properties and malformation.

| Gene | Change in geometry or physical properties | Malformations and dysfunctions |
| --- | --- | --- |
| foxp2 (2, 11) | Reduced gray matter density (thinner cortex) in Broca's area | Polymicrogyria, speech and language disorder |
| SP0535 (16) | Increased expansion of neural progenitors | improved cognitive ability and working memory |
| Cdk5 (13, 17) | Abnormalities in neuronal migration and organization | Lissencephaly with cerebellar hypoplasia (LCH) |
| GPR56 (1, 12) | Neural progenitors migration | Bilateral frontoparietal polymicrogyria |
| PAFAH1B1, RELN, TUB1A1, DCX, ARX, NDE1 (10, 19) | Increased cortical thickness | Lissencephaly |

**Table S2.** Similarity index (main text, Eq.(4)) evaluated by rescaled mean curvature of simulated and gel brain surfaces with comparison to the real brain surfaces, calculated with different vector p-norm: *p* = 1, *p* = 2, and *p* = ∞.

| Similarity Index | Simulated |  |  | Gel |  |  |
| --- | --- | --- | --- | --- | --- | --- |
| $p$ | 1 | 2 | ∞ | 1 | 2 | ∞ |
| Ferret (left) | 0.8188 | 0.7541 | 1 | 0.7448 | 0.6746 | 1 |
| Ferret (right) | 0.7733 | 0.7005 | 1 | 0.7533 | 0.6887 | 1 |
| Macaque (left) | 0.6886 | 0.6039 | 1 | 0.7658 | 0.7086 | 1 |
| Macaque (right) | 0.6832 | 0.5994 | 1 | 0.7578 | 0.6989 | 1 |
| Human (left) | 0.6056 | 0.5090 | 1 | 0.7037 | 0.6540 | 1 |
| Human (right) | 0.5911 | 0.4909 | 1 | 0.7028 | 0.6525 | 1 |

## References

Akula SK, Exposito-Alonso D, Walsh CA. Shaping the brain: The emergence of cortical structure and folding. Developmental Cell. 2023; 58(24):2836–2849.

Alenyà M, Wang X, Lefèvre J, Auzias G, Fouquet B, Eixarch E, Rousseau F, Camara O. Computational pipeline for the generation and validation of patient-specific mechanical models of brain development. Brain Multiphysics. 2022; 3:100045.

Bae BI, Tietjen I, Atabay KD, Evrony GD, Johnson MB, Asare E, Wang PP, Murayama AY, Im K, Lisgo SN, et al. Evolutionarily dynamic alternative splicing of GPR56 regulates regional cerebral cortical patterning. Science. 2014; 343(6172):764–768.

Barnette AR, Neil JJ, Kroenke CD, Griffith JL, Epstein AA, Bayly PV, Knutsen AK, Inder TE. Characterization of brain development in the ferret via MRI. Pediatric Research. 2009; 66(1):80–84.

Barresi M, Hickmott RA, Bosakhar A, Quezada S, Quigley A, Kawasaki H, Walker D, Tolcos M. Toward a better understanding of how a gyrified brain develops. Cerebral Cortex. 2024; 34(2):bhae055.

Bohi A, Wang X, Harrach M, Dinomais M, Rousseau F, Lefèvre J. Global perturbation of initial geometry in a biomechanical model of cortical morphogenesis. In: 2019 41st Annual International Conference of the IEEE Engineering in Medicine and Biology Society (EMBC) IEEE; 2019. p. 442–445.

Budday S, Raybaud C, Kuhl E. A mechanical model predicts morphological abnormalities in the developing human brain. Scientific Reports. 2014; 4(1):5644.

Bullmore ET, Bassett DS. Brain graphs: graphical models of the human brain connectome. Annual Review of Clinical Psychology. 2011; 7:113–140.

Calabrese E, Badea A, Coe CL, Lubach GR, Shi Y, Styner MA, Johnson GA. A diffusion tensor MRI atlas of the postmortem rhesus macaque brain. Neuroimage. 2015; 117:408–416.

Choi GPT, Leung-Liu Y, Gu X, Lui LM. Parallelizable global conformal parameterization of simply-connected surfaces via partial welding. SIAM Journal on Imaging Sciences. 2020; 13(3):1049–1083.

Choi GPT, Liu C, Yin S, Séjourné G, Smith RS, Walsh CA, Mahadevan L. Biophysical basis for brain folding and misfolding patterns in ferrets and humans. eLife. 2025; 14:RP107141.

Choi PT, Lam KC, Lui LM. FLASH: Fast landmark aligned spherical harmonic parameterization for genus-0 closed brain surfaces. SIAM Journal on Imaging Sciences. 2015; 8(1):67–94.

Choi PT, Lui LM. Fast disk conformal parameterization of simply-connected open surfaces. Journal of Scientific Computing. 2015; 65:1065–1090.

Cignoni P, Callieri M, Corsini M, Dellepiane M, Ganovelli F, Ranzuglia G, et al. Meshlab: an open-source mesh processing tool. In: Eurographics Italian Chapter Conference, vol. 2008 Salerno, Italy; 2008. p. 129–136.

Del-Valle-Anton L, Borrell V. Folding brains: from development to disease modeling. Physiological Reviews. 2022; 102(2):511–550.

Demirci N, Holland MA. Cortical thickness systematically varies with curvature and depth in healthy human brains. Human Brain Mapping. 2022; 43(6):2064–2084.

Garcia KE, Robinson EC, Alexopoulos D, Dierker DL, Glasser MF, Coalson TS, Ortinau CM, Rueckert D, Taber LA, Van Essen DC, et al. Dynamic patterns of cortical expansion during folding of the preterm human brain. Proceedings of the National Academy of Sciences. 2018; 115(12):3156–3161.

Gautam P, Anstey KJ, Wen W, Sachdev PS, Cherbuin N. Cortical gyrification and its relationships with cortical volume, cortical thickness, and cognitive performance in healthy mid-life adults. Behavioural Brain Research. 2015; 287:331–339.

Gill HK, Yin S, Lawlor JC, Huycke TR, Nerurkar NL, Tabin CJ, Mahadevan L. The developmental mechanics of divergent buckling patterns in the chick gut. Proceedings of the National Academy of Sciences. 2024; 121(28):e2310992121.

Gill HK, Yin S, Nerurkar NL, Lawlor JC, Lee C, Huycke TR, Mahadevan L, Tabin CJ. Hox gene activity directs physical forces to differentially shape chick small and large intestinal epithelia. Developmental Cell. 2024; 59(21):2834–2849.

Herculano-Houzel S. The human brain in numbers: a linearly scaled-up primate brain. Frontiers in Human Neuroscience. 2009; 3:857.

Heuer K, Gulban OF, Bazin PL, Osoianu A, Valabregue R, Santin M, Herbin M, Toro R. Evolution of neocortical folding: A phylogenetic comparative analysis of MRI from 34 primate species. Cortex. 2019; 118:275–291.

Hohlfeld E, Mahadevan L. Scale and nature of sulcification patterns. Physical Review Letters. 2012; 109(2):025701.

Holland MA, Miller KE, Kuhl E. Emerging brain morphologies from axonal elongation. Annals of Biomedical Engineering. 2015; 43:1640–1653.

Hu HH, Chen HY, Hung CI, Guo WY, Wu YT. Shape and curvedness analysis of brain morphology using human fetal magnetic resonance images in utero. Brain Structure and Function. 2013; 218:1451–1462.

Hutton C, Draganski B, Ashburner J, Weiskopf N. A comparison between voxel-based cortical thickness and voxel-based morphometry in normal aging. Neuroimage. 2009; 48(2):371–380.

Koenderink JJ, van Doorn AJ. Surface shape and curvature scales. Image and Vision Computing. 1992; 10(8):557–564.

Kriegeskorte N, Wei XX. Neural tuning and representational geometry. Nature Reviews Neuroscience. 2021; 22(11):703–718.

Lai CS, Gerrelli D, Monaco AP, Fisher SE, Copp AJ. FOXP2 expression during brain development coincides with adult sites of pathology in a severe speech and language disorder. Brain. 2003; 126(11):2455–2462.

Liu Z, Wang X, Newman N, Grant KA, Studholme C, Kroenke CD. Anatomical and diffusion MRI brain atlases of the fetal rhesus macaque brain at 85, 110 and 135 days gestation. Neuroimage. 2020; 206:116310.

van der Meer D, Kaufmann T. Mapping the genetic architecture of cortical morphology through neuroimaging: progress and perspectives. Translational Psychiatry. 2022; 12(1):447.

de Moraes FHP, Sudo F, Carneiro Monteiro M, de Melo BR, Mattos P, Mota B, Tovar-Moll F. Cortical folding correlates to aging and Alzheimer’s Disease’s cognitive and CSF biomarkers. Scientific Reports. 2024; 14(1):3222.

Namburete AI, Papież BW, Fernandes M, Wyburd MK, Hesse LS, Moser FA, Ismail LC, Gunier RB, Squier W, Ohuma EO, et al. Normative spatiotemporal fetal brain maturation with satisfactory development at 2 years. Nature. 2023; 623(7985):106–114.

Noctor SC, Comparisons, 2; 2016. https://ventricular.org/StephenNoctor/comparisons-2/.

Oegema R, Barakat TS, Wilke M, Stouffs K, Amrom D, Aronica E, Bahi-Buisson N, Conti V, Fry AE, Geis T, et al. International consensus recommendations on the diagnostic work-up for malformations of cortical development. Nature Reviews Neurology. 2020; 16(11):618–635.

Pang JC, Aquino KM, Oldehinkel M, Robinson PA, Fulcher BD, Breakspear M, Fornito A. Geometric constraints on human brain function. Nature. 2023; 618(7965):566–574.

Qi J, Mo F, An NA, Mi T, Wang J, Qi JT, Li X, Zhang B, Xia L, Lu Y, et al. A Human-Specific De Novo Gene Promotes Cortical Expansion and Folding. Advanced Science. 2023; 10(7):2204140.

Schneider CA, Rasband WS, Eliceiri KW. NIH Image to ImageJ: 25 years of image analysis. Nature methods. 2012; 9(7):671–675.

Schöberl J. NETGEN An advancing front 2D/3D-mesh generator based on abstract rules. Computing and visualization in science. 1997; 1(1):41–52.

Schwartz E, Nenning KH, Heuer K, Jeffery N, Bertrand OC, Toro R, Kasprian G, Prayer D, Langs G. Evolution of cortical geometry and its link to function, behaviour and ecology. Nature Communications. 2023; 14(1):2252.

Shinmyo Y, Terashita Y, Duong TAD, Horiike T, Kawasumi M, Hosomichi K, Tajima A, Kawasaki H. Folding of the cerebral cortex requires Cdk5 in upper-layer neurons in gyrencephalic mammals. Cell Reports. 2017; 20(9):2131–2143.

de Sousa AA, Beaudet A, Calvey T, Bardo A, Benoit J, Charvet CJ, Dehay C, Gómez-Robles A, Gunz P, Heuer K, et al. From fossils to mind. Communications Biology. 2023; 6(1):636.

Striedter GF, Srinivasan S, Monuki ES. Cortical folding: when, where, how, and why? Annual Review of Neuroscience. 2015; 38:291–307.

Suárez LE, Markello RD, Betzel RF, Misic B. Linking structure and function in macroscale brain networks. Trends in Cognitive Sciences. 2020; 24(4):302–315.

Takahata T, Shukla R, Yamamori T, Kaas JH. Differential expression patterns of striate cortex-enriched genes among old world, new world, and prosimian primates. Cerebral Cortex. 2012; 22(10):2313–2321.

Tallinen T, Biggins JS, Mahadevan L. Surface sulci in squeezed soft solids. Physical Review Letters. 2013; 110(2):024302.

Tallinen T, Chung JY, Biggins JS, Mahadevan L. Gyrification from constrained cortical expansion. Proceedings of the National Academy of Sciences. 2014; 111(35):12667–12672.

Tallinen T, Chung JY, Rousseau F, Girard N, Lefèvre J, Mahadevan L. On the growth and form of cortical convolutions. Nature Physics. 2016; 12(6):588–593.

Toro R, Bakker R, Delzescaux T, Evans A, Tiesinga P. FIIND: Ferret Interactive Integrated Neurodevelopment Atlas. Research Ideas and Outcomes. 2018; 4:e25312.

Toro R, Burnod Y. A morphogenetic model for the development of cortical convolutions. Cerebral cortex. 2005; 15(12):1900–1913.

Van Essen DC. A 2020 view of tension-based cortical morphogenesis. Proceedings of the National Academy of Sciences. 2020; 117(52):32868–32879.

Wang X, Lefèvre J, Bohi A, Harrach MA, Dinomais M, Rousseau F. The influence of biophysical parameters in a biomechanical model of cortical folding patterns. Scientific Reports. 2021; 11(1):7686.

Weickenmeier J. Exploring the multiphysics of the brain during development, aging, and in neurological diseases. Brain Multiphysics. 2023; 4:100068.

## References

[1] Byoung-Il Bae, Ian Tietjen, Kutay D Atabay, Gilad D Evrony, Matthew B Johnson, Ebenezer Asare, Peter P Wang, Ayako Y Murayama, Kiho Im, Steven N Lisgo, et al. Evolutionarily dynamic alternative splicing of gpr56 regulates regional cerebral cortical patterning. Science, 343(6172):764–768, 2014.

[2] Mikaela Barresi, Ryan Alexander Hickmott, Abdulhameed Bosakhar, Sebastian Quezada, Anita Quigley, Hiroshi Kawasaki, David Walker, and Mary Tolcos. Toward a better understanding of how a gyrified brain develops. Cerebral Cortex, 34(2):bhae055, 2024.

[3] Silvia Budday, Charles Raybaud, and Ellen Kuhl. A mechanical model predicts morphological abnormalities in the developing human brain. Scientific Reports, 4(1):5644, 2014.

[4] Gary P T Choi, Yusan Leung-Liu, Xianfeng Gu, and Lok Ming Lui. Parallelizable global conformal parameterization of simply-connected surfaces via partial welding. SIAM Journal on Imaging Sciences, 13(3):1049–1083, 2020.

[5] Pui Tung Choi, Ka Chun Lam, and Lok Ming Lui. Flash: Fast landmark aligned spherical harmonic parameterization for genus-0 closed brain surfaces. SIAM Journal on Imaging Sciences, 8(1):67–94, 2015.

[6] Pui Tung Choi and Lok Ming Lui. Fast disk conformal parameterization of simply-connected open surfaces. Journal of Scientific Computing, 65:1065–1090, 2015.

[7] Nagehan Demirci and Maria A Holland. Cortical thickness systematically varies with curvature and depth in healthy human brains. Human Brain Mapping, 43(6):2064–2084, 2022.

[8] Hui-Hsin Hu, Hui-Yun Chen, Chih-I Hung, Wan-Yuo Guo, and Yu-Te Wu. Shape and curvedness analysis of brain morphology using human fetal magnetic resonance images in utero. Brain Structure and Function, 218:1451–1462, 2013.

[9] Jan J Koenderink and Andrea J van Doorn. Surface shape and curvature scales. Image and Vision Computing, 10(8):557–564, 1992.

[10] Matti Koenig, William B Dobyns, and Nataliya Di Donato. Lissencephaly: update on diagnostics and clinical management. European Journal of Paediatric Neurology, 35:147–152, 2021.

[11] Cecilia SL Lai, Dianne Gerrelli, Anthony P Monaco, Simon E Fisher, and Andrew J Copp. Foxp2 expression during brain development coincides with adult sites of pathology in a severe speech and language disorder. Brain, 126(11):2455–2462, 2003.

[12] Rong Luo, Hye Min Yang, Zhaohui Jin, Dicky JJ Halley, Bernard S Chang, Lesley MacPherson, Louise Brueton, and Xianhua Piao. A novel gpr56 mutation causes bilateral frontoparietal polymicrogyria. Pediatric Neurology, 45(1):49–53, 2011.

[13] Daniella Magen, Ayala Ofir, Liron Berger, Dorit Goldsher, Ayelet Eran, Nassser Katib, Yousif Nijem, Euvgeni Vlodavsky, Shay Zur, Doron M Behar, et al. Autosomal recessive lissencephaly with cerebellar hypoplasia is associated with a loss-of-function mutation in cdk5. Human Genetics, 134:305–314, 2015.

[14] Ganeshwaran H Mochida. Genetics and biology of microcephaly and lissencephaly. Seminars in pediatric neurology, 16(3):120–126, 2009.

[15] Renske Oegema, Tahsin Stefan Barakat, Martina Wilke, Katrien Stouffs, Dina Amrom, Eleonora Aronica, Nadia Bahi-Buisson, Valerio Conti, Andrew E Fry, Tobias Geis, et al. International consensus recommendations on the diagnostic work-up for malformations of cortical development. Nature Reviews Neurology, 16(11):618–635, 2020.

[16] Jianhuan Qi, Fan Mo, Ni A An, Tingwei Mi, Jiaxin Wang, Jun-Tian Qi, Xiangshang Li, Boya Zhang, Longkuo Xia, Yingfei Lu, et al. A human-specific de novo gene promotes cortical expansion and folding. Advanced Science, 10(7):2204140, 2023.

[17] Yohei Shinmyo, Yukari Terashita, Tung Anh Dinh Duong, Toshihide Horiike, Muneo Kawasumi, Kazuyoshi Hosomichi, Atsushi Tajima, and Hiroshi Kawasaki. Folding of the cerebral cortex requires cdk5 in upper-layer neurons in gyrencephalic mammals. Cell Reports, 20(9):2131–2143, 2017.

[18] Tuomas Tallinen, Jun Young Chung, John S Biggins, and L Mahadevan. Gyrification from constrained cortical expansion. Proceedings of the National Academy of Sciences, 111(35):12667–12672, 2014.

[19] Alberto Verrotti, Alberto Spalice, Fabiana Ursitti, Laura Papetti, Rosanna Mariani, Antonella Castronovo, Mario Mastrangelo, and Paola Iannetti. New trends in neuronal migration disorders. European Journal of Paediatric Neurology, 14(1):1–12, 2010.

